# Transcriptomic balance and optimal growth are determined by cell size

**DOI:** 10.1101/2022.11.08.515578

**Authors:** Pedro J. Vidal, Alexis P. Pérez, Martí Aldea

## Abstract

Cell size and growth are intimately related across the evolutionary scale, and the molecular pathways underlying cell size homeostasis have received much attention over the last decades. However, whether cell size is important to attain maximal growth or fitness is still an open question, and the reasons why a critical size is needed for triggering key transitions of the cell cycle are unknown. We show that growth is a non-monotonic function of volume in yeast cells, with maximal values around the critical size. Comparing small to normal, large and outsized cells, the transcriptome undergoes an extensive inversion that correlates with RNA polymerase II occupancy. Accordingly, highly expressed genes impose strong negative effects on growth when their DNA/mass ratio is altered. A similar transcriptomic inversion is displayed by mouse liver cells of different sizes, suggesting that the uncovered mechanistic implications of cell size on growth and fitness are universal. We propose that cell size is set to attain a properly balanced transcriptome and, hence, maximize growth during cell proliferation.

## Introduction

Cell size is modulated by many intrinsic and extrinsic factors to comply with specific fates and developmental programs(Wood & Nurse, 2015; Ginzberg *et al*, 2015; Amodeo & Skotheim, 2016; Lloyd, 2013; Zatulovskiy & Skotheim, 2020; Kellogg & Levin, 2022; Sablowski & Gutierrez, 2022) and, among those factors, growth is perhaps the most important physiological condition. Pathways that control growth also have a vast impact in cell size (Navarro *et al*, 2012; Roberts & Lloyd, 2012; Aldea *et al*, 2017; Willis & Huang, 2017). Thus, bacteria, yeast or mammalian cells increase their mean size when they grow faster. This unambiguous phenotypic adaptation has attracted much attention and the mechanisms underlying cell size homeostasis as a function of growth have been approached in different model organisms. In turn, cells are thought to adjust their dimensions to optimize growth and fitness but the accumulated evidence is limited and the underlying causes and mechanisms are mysterious (Miettinen *et al*, 2017; Vargas-Garcia *et al*, 2018). Moreover, although it is generally accepted that cells must reach a critical size before triggering key transitions of the cell cycle, the reasons why a specific size is advantageous are also not known.

## Results

### Growth efficiency in S-G2 phases depends non-monotonically on cell size

To test whether cell size has a quantitative impact on growth, we decided to analyze the growth of single cells within a moderate range of values around the average critical size. To this end we carefully quantified the increase in volume before (G1) or after budding (S-G2) by time-lapse microscopy (Fig 1A) of cells with altered sizes by genetic ablation of activators (*CLN1, CLN2, CLN3*) or inhibitors (*WHI5, WHI3*) of Start (Johnson & Skotheim, 2013), and compared the results with the wild-type strain. The resulting linear slopes were made relative to the cell volume at the beginning of the period being analyzed to obtain the specific growth rate as an indicator of growth efficiency (Fig EV1A). We found that G1 cells were equally efficient for growth independently of their volume (Fig 1B). By contrast, the efficiency of growth in S-G2 phases followed a non-monotonic dependence on cell volume at budding (Fig 1C), the maximal efficiency being attained by wild-type cells. In agreement with these observations, cells forced to reach very large sizes during a prolonged G1 arrest exhibited a strong decrease in the specific growth rate in S-G2 phases but not in G1 (Fig EV1B). In all, cells either smaller or larger than wild-type display a decreased growth efficiency during S-G2 phases, precisely when growth rate reaches a maximum during the cell cycle in budding yeast (Ferrezuelo *et al*, 2012; Cuny *et al*, 2022) and mammalian (Tzur *et al*, 2009; Kafri *et al*, 2013) cells. The fact that this effect was not seen in G1 suggested that the critical size for triggering Start is optimized in wild-type cells to attain maximal growth rates during the posterior phases of the cell cycle. Of note, diploid and tetraploid cells did not display a decreased efficiency of growth as a function of cell volume (Fig 1D).

**Figure 1.**
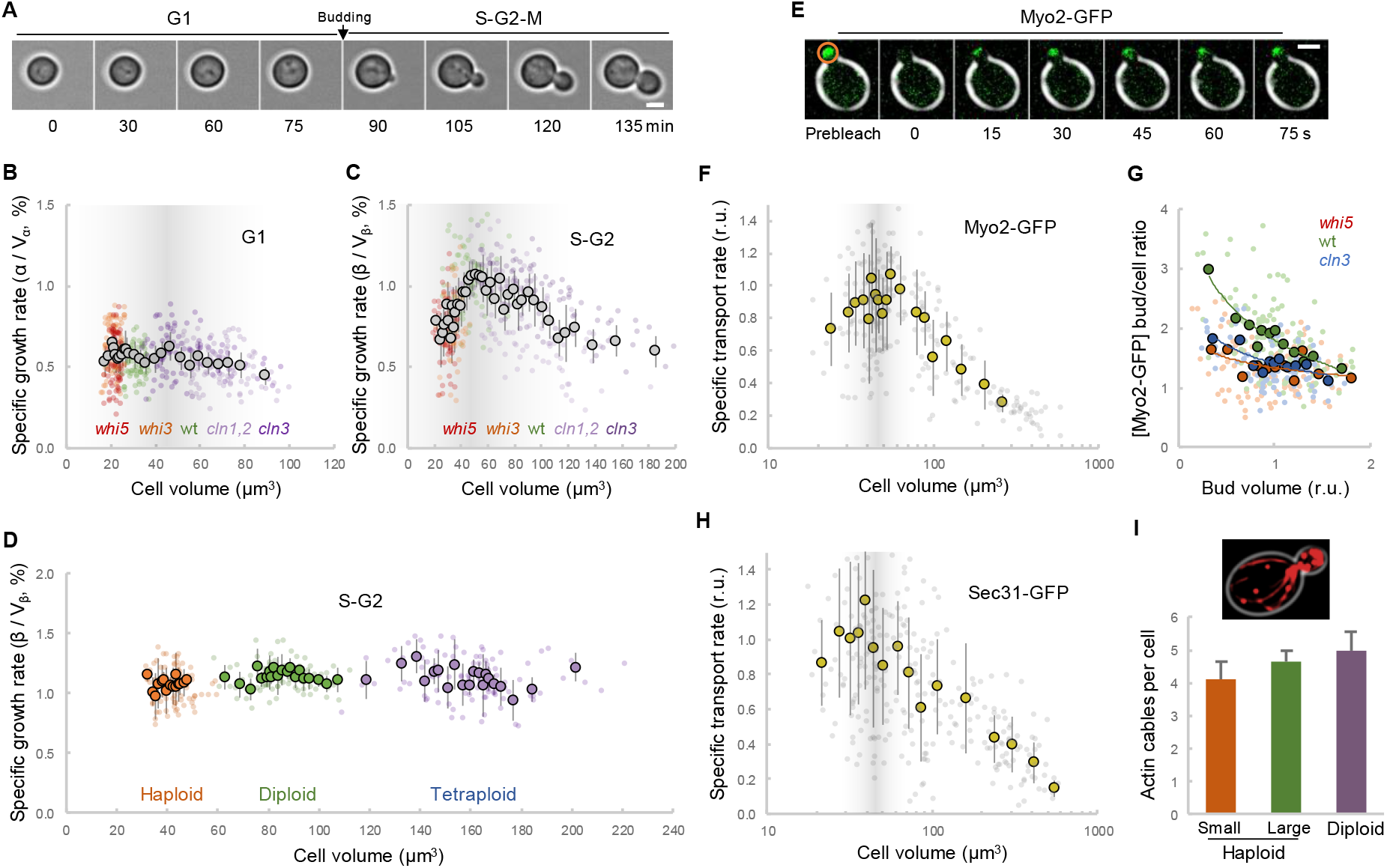
Growth potential in S-G2 phases is maximal around the critical size. A A representative yeast cell growing during the cell cycle. Scale bar, 2 µm. B-D Specific growth rates from G1 (B) and S-G2 (C,D) cells with the indicated genotypes as a function of cell volume. Single-cell data (N = 100 for each genotype, small circles) are plotted. Mean values of binned data (bin size = 20, large circles) with confidence limits for the mean (α = 0.05) are also shown. The diffuse gray bar in B and C represents the distribution of critical sizes from wild-type (wt) cells. E FRAP analysis of Myo2-GFP in a representative yeast cell shortly after budding. Scale bar, 2 µm. F Specific transport rates of Myo2-GFP as a function of cell volume from cells as in B and Fig EV1B. Relative single-cell data (N = 220, small circles) are plotted. Mean values of binned data (bin size = 10, large circles) with confidence limits for the mean (α = 0.05) are also shown. The diffuse gray bar represents the distribution of critical sizes from wild-type cells. G Bud to cell ratios in Myo2 concentration as a function of bud volume from cells with the indicated genotypes. Single-cell data (N = 100 for each genotype, small circles) are plotted. Mean values of binned data (bin size = 10, large circles) with confidence limits for the mean (α = 0.05) and the corresponding logarithmic regression lines are also shown. H Specific transport rates of Sec31-GFP as a function of cell volume as in F. Relative single-cell data (N = 284, small circles) are plotted. Mean values of binned data (bin size = 10, large circles) with confidence limits for the mean (α = 0.05) are also shown. The diffuse gray bar represents the distribution of critical sizes from wild-type cells. I Number of actin cables in small and large haploid cells compared to diploid cells. Mean values (N = 15) per cell and confidence limits for the mean (α = 0.05) are plotted. Inset: A representative cell expressing Abp140-mCherry to stain the actin cytoskeleton is shown.

Growth in S-G2 phases is polarized and requires the actin-cytoskeleton transport machinery to drive the accumulation of new cellular components to the bud. Thus, we decided to test whether polarized transport depends on cell volume by analyzing Myo2, a key motor protein involved in actin-based transport of vesicular cargos. We used fluorescence recovery after photobleaching (FRAP) in the bud compartment of small budded S-G2 cells (Fig 1E) and, as expected, Myo2-GFP recovery was strongly reduced by an actin depolymerization agent (Fig EV1C and D). As discussed above, the input rate in each bud was made relative to the volume of the mother cell compartment to obtain the specific rate as an indicator of transport efficiency. We found that the specific transport rate was maximal in wild-type cells, and decreased in both smaller and larger cells (Fig 1F) as observed for growth efficiency. Giving support to the notion that active transport of Myo2 to the bud is maximal in wild-type cells, the ratio of steady-state concentrations in the bud and mother cell compartments was also higher in wild-type cells compared to either small (*whi5*) or large (*cln3*) cells (Fig 1G), this effect being more pronounced in small buds, in which the relative effect of the input rate should be more important. Sec31, a component of the COPII vesicle coat, displayed a very similar behavior, with the highest specific transport rates in wild-type cells (Figs 1H and EV1D). Whereas the concentration of Myo2 was nearly constant over a large range of cell volumes (Fig EV1E), the number of actin cables per cell only increased by 30% over a 10-fold cell volume increase (Fig EV1F). However, actin-cable number in diploid and large haploid cells was not significantly different (Fig 1I). It is worth noting that the diameter of the bud neck has been shown to scale with cell size (Kukhtevich *et al*, 2020). In all, these data would rule out a limiting effect of the actin cytoskeleton on polarized transport through the bud neck at large cell sizes. In contrast, considering that tetraploid and diploid cells did not display altered specific growth rates, our results point to the notion that DNA could become limiting, likely as template for transcription, during S-G2 phases in large haploid cells.

### The transcriptome becomes widely perturbed in outsized cells

To deepen into the defects caused by size in the specific growth rate of S-G2 cells we compared the transcriptomes of control and outsized cells shortly after budding. To obtain these two populations, cells conditionally expressing G1-cyclin (*GAL1p-CLN3 cln1,2*) in galactose were shifted to glucose for 90 min to synchronize them in G1, or 8 h to increase cell size during the prolonged G1 arrest, and subsequently transferred to a galactose-based medium to induce *CLN3* expression, trigger cell cycle entry, and reach a 50% budding frequency. In this manner we obtained synchronous cells in S-G2 phases with average volumes of either 53 fl (*CLN3*^ON^, control cells) or 456 fl (*CLN3*^OFF>ON^, outsized cells) that were immediately used to extract total RNA for transcriptomic analysis. Expression of more than 6000 genes was consistently detected in three independent experiments. Remarkably, 45.8% of them displayed significant differences (FDR < 0.05) in relative mRNA levels; 2058 genes were upregulated and 718 genes were downregulated (Fig 2A), indicating that almost half of the transcriptome becomes perturbed when cells enter the cell cycle with a very large size.

**Figure 2.**
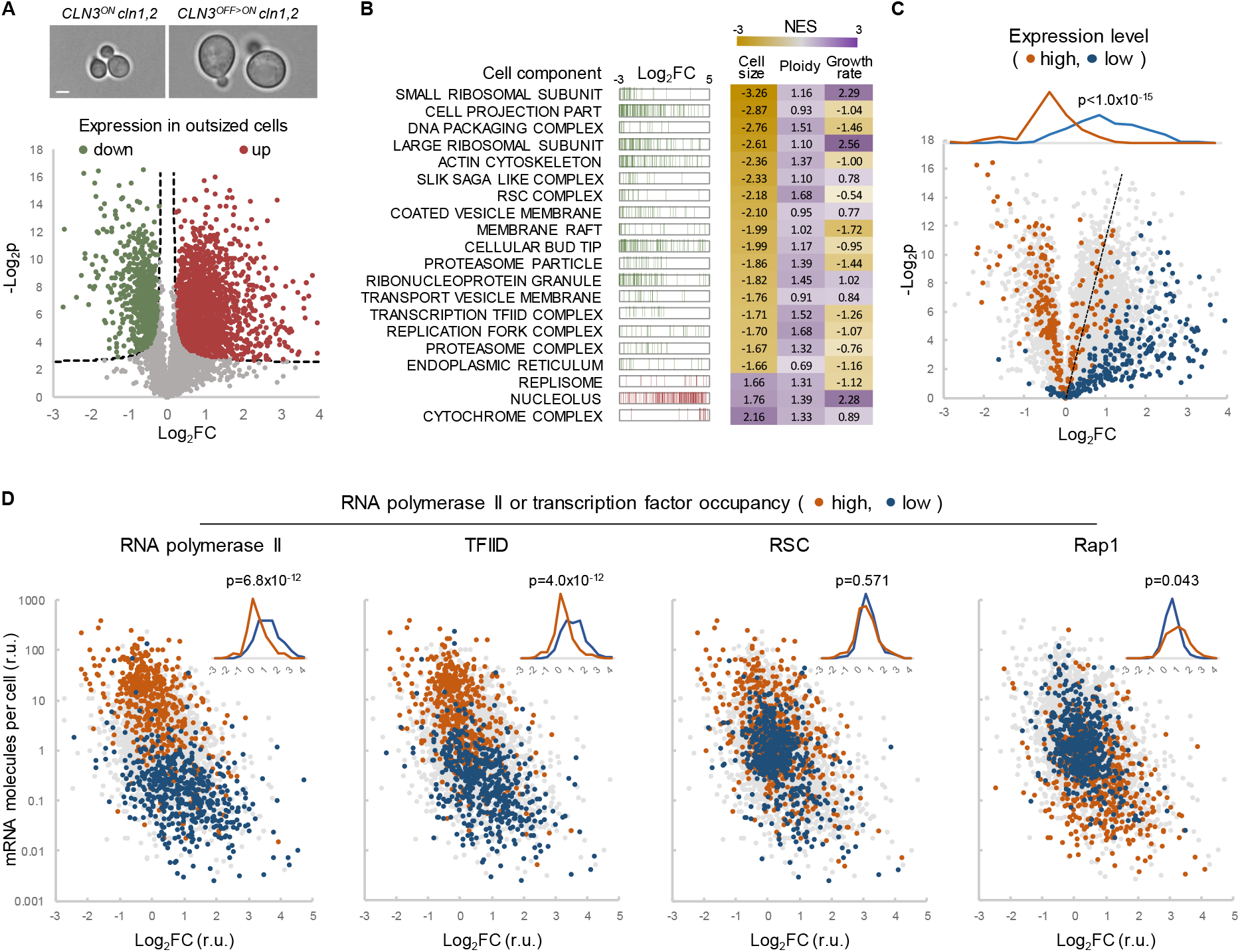
The transcriptome is extensively imbalanced in outsized cells. A Volcano plot comparing the transcriptomes of outsized and control cells. G1-cyclin conditional expression was used to synchronize (control, *CLN3*^*ON*^ *cln1,2*) or arrest cells in G1 for 8h (outsized, *CLN3*^*OFF>ON*^ *cln1,2*) and induce them to enter the cell cycle by turning on *CLN3* expression. Representative cells are shown. Scale bar, 2 µm. The plot shows t-test significance (-Log_2_p) as a function of fold change (Log2FC) in expression of yeast genes (N = 6059, gray) comparing outsized to control cells. Significantly (FDR < 0.05) downregulated (N = 718, green) or upregulated (N = 2058, red) genes are highlighted. B Gene enrichment analysis of the transcriptomic imbalance in outsized cells. Barcode plots indicate the rank position of genes from the corresponding GO cell component that were found significantly (FDR<0.05) enriched within downregulated (green) or upregulated (red) genes in outsized cells. Normalized enrichment scores (NES) are shown at the right with those obtained from the ploidy and growth rate datasets for the same GO cell components. C Volcano plot as in A highlighting genes that are expressed at high (top 200 genes, orange) or low (bottom 200 genes, blue) levels in wild-type cells. The dotted line splits the dataset into two equal moieties. Inset shows their frequency distributions as a function of the expression fold change in outsized cells with the result of a Mann-Whitney U test. D Plots of wild-type mRNA levels as a function of the expression fold change (Log_2_FC) in outsized cells highlighting genes with high (top 500 genes, orange) or low (bottom 500 genes, blue) occupancy of RNA polymerase II or the indicated transcription factors in wild-type cells. Insets show the corresponding frequency distributions as a function of the expression fold change in outsized cells with the result of a Mann-Whitney U test.

Genes downregulated in outsized cells were enriched in several Gene Ontology (GO) terms for cell components (Figs 2B and EV2A), including ribosomes, the polar growth machinery, nucleosomes, DNA-remodeling complexes, the proteasome and the TFIID complex. By contrast, none of these cell components were enriched among downregulated genes in large tetraploid cells (Fig 2B) as deduced from the analysis of existing datasets (Wu *et al*, 2010). In addition, contrary to the behavior we observed, ribosomal protein genes were upregulated in large fast-growing cells compared to small nutrient-limited cells (Slavov & Botstein, 2011). The results of these analyses point to the notion that the relative downregulation of genes belonging to the abovementioned cell components in outsized haploid cells is not due to a larger cell size *per se* but to an inherently altered mass/ploidy ratio. Similarly to histones, inhibitors of Start were downregulated in outsized cells (Fig EV2B) as found by others (Chen *et al*, 2020; Swaffer *et al*, 2021a). Genes upregulated in outsized cells belonged to a wide panoply of cell components, with only the cytochrome complex and the nucleolus offering a clear enrichment that was maintained in large tetraploid and fast-growing cells (Figs 2B and EV2A).

### The transcriptomic imbalance in outsized cells is a function of expression levels and pre-initiation complex occupancy

Although some of the genes coding for the TFIID complex were downregulated in outsized cells, we found that transcript levels of all other components of the pre-initiation complex (PIC) were relatively constant in outsized cells (Fig EV2B) as observed by others (Swaffer *et al*, 2021a). In addition, the relative decrease of TFIID components (log_2_FC=-0.08±0.22) was very small compared to histones (log_2_FC=-1.16±0.61), which have been shown to be maintained constant relative to ploidy (Wu *et al*, 2010). In all, the effective PIC/DNA ratio should be expected to increase quasi-linearly with cell size, which might differently affect gene expression as a function of the relative affinity of every promoter for the PIC and, hence, of the corresponding transcriptional output. In support of this idea, we found that high expression genes were mostly downregulated in outsized cells while low expression genes were typically upregulated (Fig 2C). Moreover, when all genes were considered, we obtained a strong negative correlation between gene expression in control cells and the fold change in outsized *vs*. control cells (Fig EV2C). Accordingly, the enrichment scores of high and low expression genes within the transcriptomic dataset from outsized cells were strongly negative and positive, respectively (Fig EV2D). Notably, no enrichments of high and low expression genes were detected in the transcriptomic dataset from large tetraploid cells, which would support the idea that the observed transcriptional imbalance is due to the altered PIC/ploidy ratio inherent to outsized cells. We also obtained the enrichment scores within the dataset from large fast-growing cells, but they were of opposite sign to those for the outsized cells dataset (Fig EV2D). As expected, highly-expressed genes such as those coding for ribosomal proteins are upregulated when cells grow faster (Slavov & Botstein, 2011).

The ribosome was the cell component whose encoding genes displayed the most negative enrichment score within the transcriptomic dataset from outsized cells (Fig EV2E). Surprisingly, ribosome biogenesis genes behaved in a completely opposite manner and offered a positive enrichment score, which suggests that the imbalance in the synthesis of ribosomal proteins is not the result of a concerted physiological response to an increase in cell size. Moreover, since Hsf1-dependent genes of the environmental stress response (ESR) were not upregulated in outsized cells, a role of this pathway in downregulating ribosomal protein expression (Albert *et al*, 2019) would be unlikely.

As expected from their extremely high expression levels, ribosomal protein genes are particularly enriched in RNA polymerase II and TFIID (Fig EV2F), suggesting that PIC availability could be involved in the expression imbalance of these genes in outsized cells. Thus, we decided to compare available datasets (Rossi *et al*, 2021) on gene occupancy by different components of the PIC with the fold change in gene expression in outsized *vs*. control cells. Genes displaying high occupancy by Rpb1 (the major subunit of RNA polymerase II), Spt15 (the TATA-binding protein of TFIID) and Sua7 (TFIIB) were mostly downregulated, while genes with low occupancy were clearly upregulated (Figs 2D and EV2G). By contrast, occupancy by components of SAGA and RSC, two chromatin remodelers, were not correlated with gene expression fold change in outsized cells.

Finally, gene occupancy by specific transcription factors important for growth as Rap1, Fhl1 or Ifh1 displayed a small, albeit significant correlation with gene expression fold change in outsized cells, but of opposite sign to PIC occupancy (Figs 2D and EV2G). In all, these results suggest that an increased PIC/ploidy ratio and the diverse promoter affinities for the PIC would underlie the extensive perturbation of gene expression in outsized cells.

### Many highly-expressed genes are haploinsufficient and limiting for growth in S-G2

To address the functional relevance of the transcriptomic imbalance of highly-expressed genes in outsized cells, we first compared available data on haploinsufficiency at a genomic level (Pir *et al*, 2012) with the expression fold change in outsized *vs*. control cells. The most haploinsufficient genes showed a negative enrichment score within the transcriptomic dataset from outsized cells (Fig 3A), i.e. their expression was mostly downregulated at very large cell sizes. By contrast, haploproficient genes were not significantly enriched. We then plotted the growth-rate change from the genomic haploinsufficiency study as a function of expression levels or RNA polymerase II occupancy in wild-type cells, and found that highly-expressed or densely RNA polymerase II-occupied genes displayed a clear decrease in growth efficiency of the corresponding hemizygous diploid cells (Fig EV3A and B). In other words, many highly-expressed genes become limiting for growth if their copy number is reduced.

**Figure 3.**
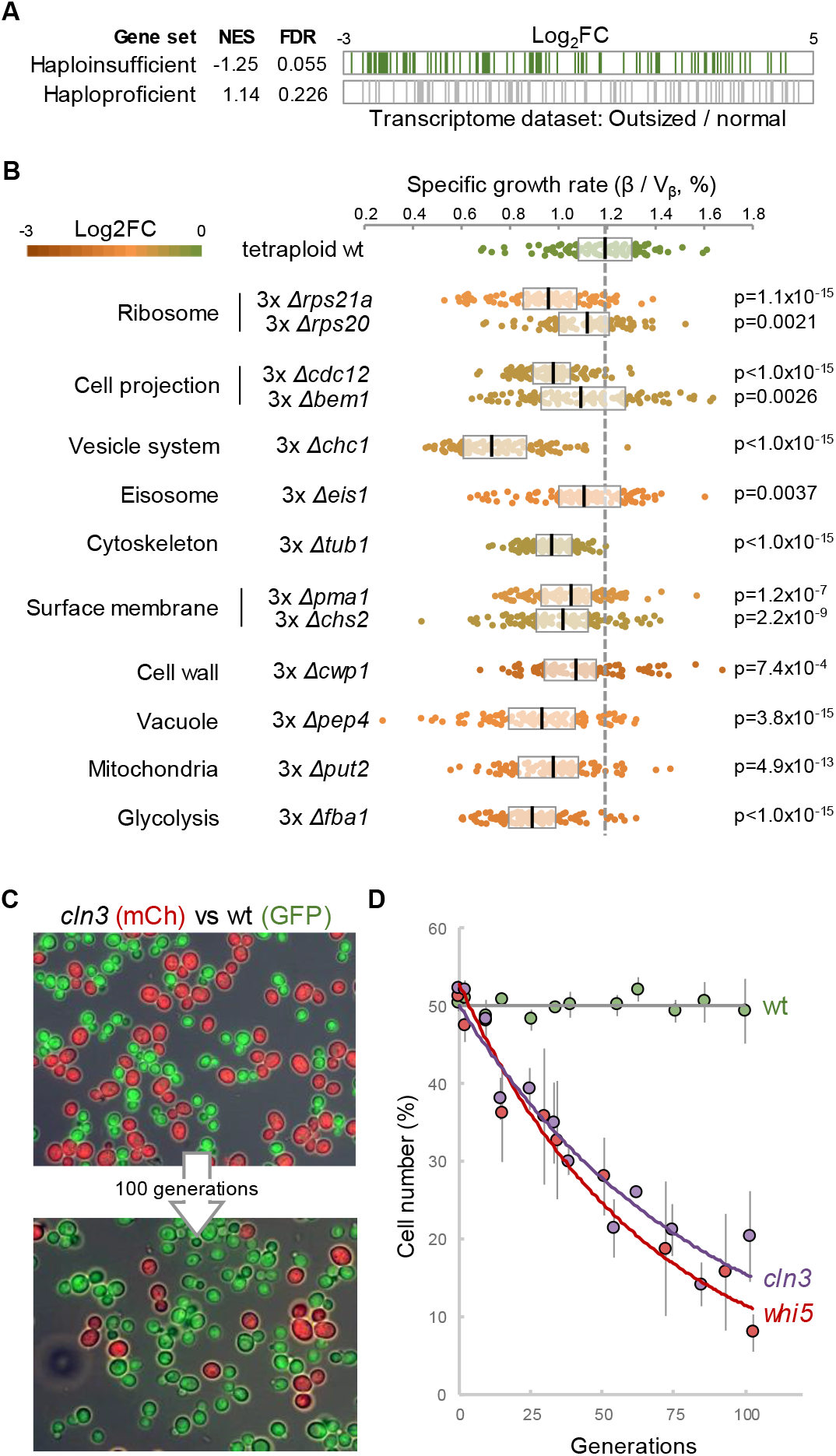
Cell size, growth competence and haploinsufficiency. A Gene enrichment analysis of haplo-insufficiency within the transcriptomic dataset of outsized *vs*. control cells. Barcode plots show the position of genes displaying strong haploinsufficiency (N = 100) or haploproficiency (N = 100). Normalized enrichment scores (NES) and the corresponding FDR values are indicated. B Specific growth rates from S-G2 cells with the indicated genotypes. Single-cell data (N = 100 for each genotype, small circles) are plotted with the corresponding median and quartile values. Pairwise comparisons with the wild-type were performed with a Mann-Whitney U test, and the resulting p-values are indicated. C Competition experiments of cells with different sizes. Representative images of initially equal numbers of large (*cln3*, red) mCherry-expressing cells and normal GFP-expressing (wild-type, green) cells grown for 100 generations by consecutive dilution. D Cell frequencies of small (*whi5*, red), normal (wild-type, green), or large (*cln3*, purple) mCherry-expressing cells competing with wild-type GFP-expressing cells as a function of generation number. Mean ± se values are plotted.

Next, we wanted to know if downregulated genes in outsized cells have a copy-number dependent effect on cell fitness. To this end we chose 13 representative genes of different cell components among those strongly downregulated in outsized cells, and built tetraploid strains in which three copies of the corresponding gene had been deleted. After analyzing these strains by time-lapse microscopy, we found that copy-number reduction of any of these genes produced a significant decrease (5% to 40%) in the specific growth rate in S-G2 (Fig 3B), indicating that expression of these genes is limiting for maximal growth.

Finally, to ascertain if altered mass/ploidy ratios have an impact on cell fitness, we mixed equal numbers of small (*whi5*), normal (wild-type), or large (*cln3*) cells expressing mCherry and wild-type cells expressing GFP, and grew them for many generations by consecutive dilution of batch cultures (Fig 3C). We found that both small and large cells were readily outcompeted by wild-type cells and reduced their presence to *ca*. 10% in 100 generations (Fig 3D).

### The transcriptome is inverted according to cell size

The results described so far suggested that cell size, by affecting the PIC/ploidy ratio, could progressively alter the relative contribution of every gene to the transcriptome as a function of its affinity for the PIC and, hence, RNA polymerase II occupancy. Thus, we decided to analyze the transcriptomes of small (*whi5*), normal (wild-type) and large (*cln3*) cells, and perform a comparative analysis with outsized (*CLN3*^OFF>ON^ *cln1,2*) cells (Fig 4A and B). The total mRNA amount per cell volume was rather constant from small (*whi5*) to large (cln3) cells, but it dropped by 60% in outsized cells (Fig 4C) as previously described (Neurohr *et al*, 2019). Regarding gene expression dependence on cell size, we first obtained the fold change from large (*cln3*) to small (*whi5*) cells and compared the resulting values to those obtained from outsized (*CLN3*^OFF>ON^ *cln1,2*) and control (*CLN3*^ON^ *cln1,2*) cells. As observed in Fig EV4A, there was a good correlation along the full range of fold change values between the two experimental datasets, indicating that the overall transcriptomic effects observed when comparing control (52±17 fl) and outsized (550±260 fl) S-G2 cells are also observed when comparing small (31±8 fl) and large (88±24 fl) cycling cells within a range of cell volumes closer to the wild-type (40±9 fl) strain. More important, as in the experiments with outsized cells, we found that high and low expression genes were downregulated and upregulated, respectively, when comparing large (*cln3*) to small (*whi5*) cells (Fig EV4B). Also, ribosomal and nucleolar protein genes displayed the same opposite behavior as in the outsized *vs*. control datasets (Fig EV4C).

**Figure 4.**
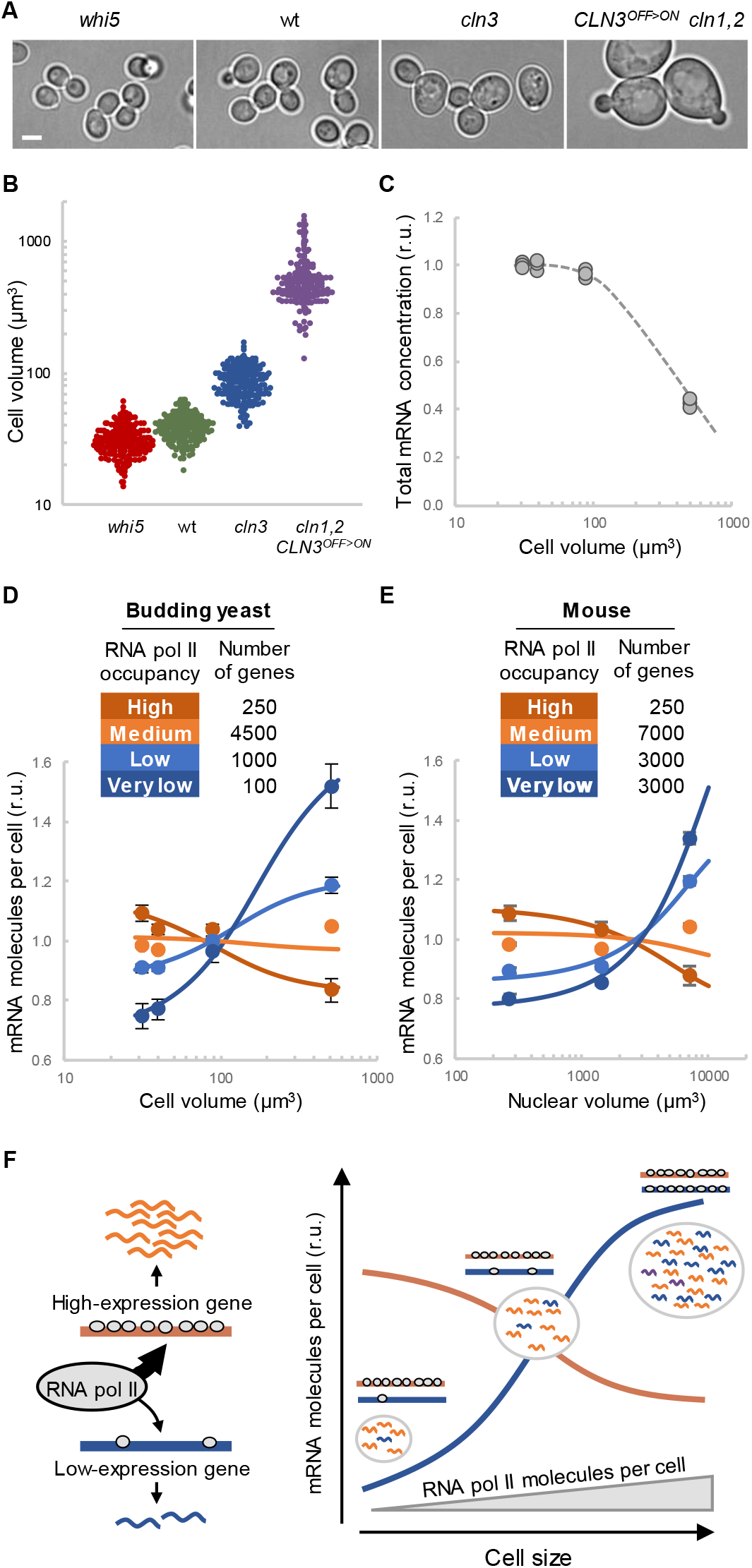
Inversion of the transcriptome by cell size and RNA polymerase II occupancy. A,B Small (*whi5*), normal (wild-type, wt), large (*cln3*) and outsized (*CLN3*^OFF>ON^ *cln1,2*) cells used for transcriptomic analysis. Representative images (A) and a plot with single-cell (N = 200) volume data (B) are shown. Scale bar, 2 µm. C Total mRNA concentration as a function of cell volume. Relative data from three independent datasets are shown with a regression line. D,E Relative expression levels of yeast (D) and mouse (E) genes grouped by high, medium, low and very low RNA polymerase II occupancy as a function of cell or nuclear volume, respectively. Mean values and confidence limits (α = 0.05) are plotted. Lines were obtained by fitting the model to the corresponding experimental data. F Basic elements of the model used to recapitulate the transcriptomic inversion as a function of cell size. First, high and low expression genes have different affinities for the PIC and occupancy by RNA polymerase II. Second, the number of RNA polymerase II and PIC molecules per cell increases with cell size. As RNA polymerase II is a key limiting factor for transcription, high (strongly saturated) and low (not saturated) expression genes respond with different dynamics leading to a transcriptional inversion.

Next we ranked genes by RNA polymerase II occupancy and group them in four categories: high, medium, low and very low occupancy. To facilitate comparison of the relative changes within each dataset, mRNA levels (rpkm) obtained from every gene were normalized to the average obtained from the corresponding gene at the four different cell volumes analyzed. Notably, while genes with high RNA polymerase II occupancy were relatively downregulated as a function of cell size from small to wild-type, large and outsized cells, genes with very low RNA polymerase II occupancy showed a strong relative upregulation with increasing cell size (Fig 4D). Genes with medium and low RNA polymerase II occupancy displayed an intermediate behavior, causing a general and progressive inversion of the transcriptome in relative terms as a function of cell size from small to outsized cells. Expression differences of genes with high and very low RNA polymerase II occupancy in small (*whi5*) and wild-type cells were not outstanding but significant (p=9.3×10^−7^ and p=0.05, respectively). Gene expression and RNA polymerase II occupancy have been analyzed in mouse liver cells of different sizes by conditional CDK1 expression during proliferation after tissue resection (Miettinen *et al*, 2014). Notably, when we reanalyzed these datasets grouping genes by RNA polymerase II occupancy we found a similar inversion in relative mRNA levels as a function of cell size (Fig 4E).

Finally, we wanted to know whether a simple model based on different affinities for the transcriptional machinery and increasing RNA polymerase II/DNA ratios as cells become large (Fig 4F) would recapitulate the observed transcriptomic inversion in gene expression with cell size. To this end, we simulated expression of four gene sets as in Fig 4D with realistic nuclear volumes and RNA polymerase II and promoter DNA concentrations. First, we found that the RNA polymerase II binding constant was the key parameter modifying the pattern of expression dependence on cell size (Fig EV4D). After fitting the RNA polymerase II binding constants of the four gene sets to the experimental dataset, the model produced a very similar transcriptomic inversion within the experimental range of cell sizes (see lines in Fig 4D). Of note, the fitted binding constants (10^6^ to 10^7^ M^−1^) were very close to those experimentally estimated for TFIIB (2×10^7^ M^−1^)(Lagrange *et al*, 1998). Regarding the overall RNA polymerase II concentration, the fitted value (6×10^−7^ M) agrees well with available experimental data (Ho *et al*, 2018). In addition, the fitted model predicted a decrease in the overall mRNA concentration at very large cell sizes as experimentally observed (Fig EV4E). The cell-volume range in which the model causes the transcriptomic inversion coincides with the range wherein the absolute number of RNA polymerase II molecules overcomes the number of gene promoters (Fig EV4D). In this regard, by reanalyzing a transcriptomic dataset of the effects of a HDAC inhibitor that derepresses poorly-expressed genes (Tirosh *et al*, 2009), we observed a clear upregulation of genes with high RNA polymerase II occupancy (Fig EV4F), which supports the general limiting role of RNA polymerase II and other PIC components in genome wide transcription. Last, the model was also able to recapitulate the transcriptomic inversion observed in mouse liver cells as a function of nuclear volume (see lines in Fig 4E). The predicted numbers of RNA polymerase II molecules as a function of nuclear volume were lower than experimentally observed, suggesting that parameters of the transcription process other than RNA polymerase II binding would be limiting in mammalian cells. In all, these results support the notion that intrinsic differential affinity for the PIC causes distinct dependence patterns of gene expression on cell volume, thus providing a mechanistic framework to set growth as a non-monotonic function of cell size.

## Discussion

We demonstrate that growth rate in S-G2 phases is a non-monotonic function of cell volume, with the highest values within a narrow range around the critical size of wild-type cells. Growth-dependence on cell size was lost at constant mass/ploidy ratios, pointing to gene dosage as the key factor limiting protein synthesis and growth in S-G2 phases. We found that almost half of the transcriptome becomes perturbed in outsized cells. Gene deregulation displayed a strong correlation with gene expression and occupancy by RNA polymerase II, TFIID and TFIIB, but not by chromatin remodeling factors or specific transcription factors deeply involved in growth regulation. By analyzing a range of volumes around the critical size, we found that expression of genes grouped by RNA polymerase II occupancy progressively decreases when comparing small to normal, large and outsized cells, leading to a transcriptomic inversion along cell size.

Genes have different affinities for the transcriptional machinery to modulate their expression and, in turn, efficiently coordinate all cellular processes. In other words, cells need gene expression to be precisely balanced at a genome-wide level to maximize cell fitness. In the case of budding yeast, we have demonstrated the importance of reaching a critical size at the G1/S transition to attain the highest growth rate in the posterior S-G2 phases of the cell cycle. Therefore, in proliferating cells, the critical size would emerge as the volume in which the proper transcriptomic equilibrium is attained to maximize growth. In smaller and larger cells, the transcriptomic balance is tilted in opposite directions thus perturbing the coordination of gene expression at a genome wide level which, in either case, would necessarily have a negative impact on cell fitness. Several independent lines of evidence give support to this hypothesis. First, we have found that most genes with a strong haploinsufficiency score are downregulated in large cells. Also, highly-expressed or densely RNA polymerase II-occupied genes display a clear decrease in growth efficiency when placed in hemizygosis. Third, a relevant sample of high-expression genes that are downregulated in large cells displays a strong reduction in growth efficiency in S-G2 phases when expressed from a single copy in tetraploid cells. Finally, we also show that both small and large cells display a decreased fitness when forced to compete with wild-type cells during sustained proliferation. In all, we conclude that specific affinities of core promoters for the PIC and the PIC/DNA ratio, which is a function of cell size, are optimized in wild-type cells to attain maximal growth in S-G2 phases. These relationships underscore the importance of a cell-size checkpoint at Start, shortly before progression into S-G2-M phases, when yeast and mammalian cells reach the highest rates of growth (Cuny *et al*, 2022; Ferrezuelo *et al*, 2012; Kafri *et al*, 2013; Tzur *et al*, 2009) and carbohydrate oxidative metabolism (Burnetti *et al*, 2016; Papagiannakis *et al*, 2016; Wang *et al*, 2014; Zhao *et al*, 2016; Ewald *et al*, 2016) within the cell cycle to fuel DNA replication and mitosis.

Our data fully support a recent theoretical study on the dependence of gene expression on cell size (Wang & Lin, 2021), which predicted that genes with high or low RNA polymerase II recruitment capacity would be relatively downregulated or upregulated, respectively, in large cells. Based on gene enrichment analysis of a published dataset (Chen *et al*, 2020), the authors related their findings to the possible limiting nature of specific transcription factors. However, a recent study has shown that RNA polymerase II is a major limiting component of the transcriptional machinery (Swaffer *et al*, 2021b), suggesting that genes compete for finite PIC numbers. Taking into account that the PIC/DNA ratio increases with cell size, we propose a general mechanism in which differential PIC recruitment dynamics is the key factor causing many genes to display non-linear scaling of expression with cell size. In this regard, a genome-wide screen for size-controlling factors concluded that genetic defects in RNA polymerase II and general transcription factors compromise cell size homeostasis (Maitra *et al*, 2019). Gene occupancy by remodeling factors did not correlate with the gene expression fold change in outsized *vs*. control cells, which supports the finding that chromatin accessibility is not modulated by cell size (Swaffer *et al*, 2021b). Although we cannot rule out that other steps of the gene expression process are modulated by cell dimensions, mRNA scaling by cell size has been associated to transcription initiation, but not to mRNA decay, in fission yeast (Sun *et al*, 2020).

If the PIC/DNA ratio and, hence, cell size at Start are set to attain a properly balanced transcriptome, the inverse relationship should also be considered. Chen et al. (Chen *et al*, 2020) have shown that most activators of Start are upregulated in large G1 cells, whereas most inhibitors display the opposite behavior and become downregulated when cells become larger, and propose that the ratio of activators to inhibitors triggers cell-cycle entry. Due to the lack of correlation with a ploidy dataset, these authors suggest that scaling in gene expression is due to the mass/DNA ratio, rather than size *per se*, as we find in this work. Thus, differential scaling of activator and inhibitory gene expression with cell size could be due to differences in their affinity for the PIC components. However, activator and inhibitory genes of Start do not display clear differences in RNA polymerase II occupancy, pointing to the existence of other mechanisms driving non-linear scaling of gene expression with cell size.

As shown by others in yeast (Swaffer *et al*, 2021b; Zhurinsky *et al*, 2010; Sun *et al*, 2020) and mammalian cells (Padovan-Merhar *et al*, 2015), we found that expression of most genes largely scales with cell size within a physiological range, but not in very large cells, were we observed a strong decrease in the overall concentration of mRNAs as previously found (Neurohr *et al*, 2019). A theoretical approach to understand transcriptional homeostasis during growth as cells progress through the cell cycle predicted that DNA would limit gene expression in very large haploid cells (Lin & Amir, 2018). However, we found that mRNAs coding for PIC components decreased their concentration in very large cells like most other mRNAs, suggesting that the low biosynthetic capacity of very large cells is due to the combination of multiple effects.

A non-monotonic dependency of growth on cell size was recently proposed as a mechanism to maintain cell dimensions within limits during growth (Barber *et al*, 2020). If small and large cells grew with lower efficiencies, they would replicate more slowly and be readily counterselected in a growing population. Although our competition experiments support these ideas, differences in fitness are only observed after 20 generations and would only explain a fraction of cell size control across consecutive divisions.

Mammalian cells display a similar non-monotonic dependence of fitness on cell size (Miettinen & Björklund, 2016). In this regard, here we show that a transcriptomic inversion also occurs in mouse hepatocytes of different sizes induced to proliferate after liver resection (Miettinen *et al*, 2014). These observations suggest that the mechanistic framework that maximizes growth and/or fitness as a function of cell size is universal. As a striking example, zygotic transcription in the early embryo is inhibited until initial divisions reduce cell size below a precise threshold (Chen *et al*, 2019; Jukam *et al*, 2021). We envisage that this critical size is set to generate a properly balanced transcriptome and, hence, efficient execution of the subsequent phases of embryonic development. The same strategy could also apply to other circumstances (Hubatsch *et al*, 2019), wherein modulation of cell size would provide the specifically suited balance in gene expression at a genome wide level.

## Materials and Methods

### Strain constructions

Yeast strains used in this study are listed in Table EV1. Methods for chromosomal gene transplacement and PCR-based directed mutagenesis have been described (Ferrezuelo *et al*, 2012). Unless stated otherwise, all gene fusions were expressed at endogenous levels at their respective loci. Gene fusions and specific constructs requiring multiple fragments were obtained by one-step recombination in yeast cells as described (Gibson *et al*, 2008). Tetraploid **a**/**a**/**α**/**α** cells were originated from diploid cells following published procedures (Xie *et al*, 2017).

### Growth conditions

Cells were grown in SC medium with 2% glucose or 2% galactose at 30°C unless stated otherwise. Where indicated, latrunculin A was added at 100 μM.

### Live cell microscopy

Cells were imaged by time-lapse microscopy in 35-mm glass-bottom culture dishes (GWST-3522, WillCo) in SC-based media at 30°C essentially as described (Ferrezuelo *et al*, 2012) using fully-motorized Leica AF7000 or Thunder Imager microscopes. Time-lapse images were analyzed with the aid of BudJ (Ferrezuelo *et al*, 2012), an ImageJ (Wayne Rasband, NIH) plugin (https://www.ibmb.csic.es/en/department-of-cells-and-tissues/spatial-control-of-cell-cycle-entry/#lab-corner) to obtain cell dimensions and fluorescence data. Volume growth rate in G1 and S-G2 was determined as described in Fig EV1A.

### Transport rate determinations by FRAP

To analyze transport kinetics of Myo2-GFP and Sec31-GFP, a circle circumscribing the bud was photobleached at 488-nm until fluorescence dropped to ca. 5% of the initial value, and the bud was imaged every 1 sec to record fluorescence recovery under a Zeiss LSM780 confocal microscope. After background subtraction, fluorescence data were corrected with those from a non-bleached cell, and the total fluorescence in the bud compartment during recovery was used to fit a logarithmic function. The resulting slope was divided by the mother cell volume to obtain the specific transport rate.

### RNA extraction, sequencing and data analysis

Before RNA extraction, culture samples were added a fixed number of *Pichia pastoris* cells (*ca*. 1:100) to allow calculations of comparable relative mRNA molecule numbers per cell and relative concentrations. Total RNA was obtained by a phenol-based extraction method(Gallego *et al*, 1997), and used to prepare polyA^+^ enriched libraries for paired-end (2×100 bp) sequencing (BGI DNBSEQ platform). More than 30 million reads were sequenced per sample, which were aligned to the *S. cerevisiae* and *P. pastoris* genomes using HISAT2 (Kim *et al*, 2015) in the Galaxy platform (Boekel *et al*, 2015). Assignment to transcriptional units, standard filtering steps and basic statistical analysis of triplicate samples was done using CoverageView (version 1.32.0, Lowy, E., 2021) in Bioconductor (Huber *et al*, 2015) and custom R scripts. Due to the specific genetic traits for G1-cyclin conditional expression, outsized (*CLN3*^OFF>ON^) and control (*CLN3*^ON^) cells were grown under different conditions compared to small (*whi5*), normal (wild-type) and large (*cln3*) cells. To compare the two transcriptomic datasets, rpkm data from outsized cells were multiplied by the corresponding wild-type/control rpkm ratio. **ChIP-seq data analysis**. Gene occupancy by Rpb1 (RNA polymerase II), Spt15 (TFIID), Sua7 (TFIIB), Spt20 (SAGA) and Snf2 (RSC) was analyzed from available ChIP-exo datasets in project PRJNA622558 (Rossi *et al*, 2021). Promoter binding by Rap1, Ifh1, and Fhl1 in the yeast genome was analyzed from available ChIP datasets in project PRJNA261651 (Knight *et al*, 2014). Read alignment and assignment procedures were as above with the following specific conditions: for Rpb1 coverage, reads matching only the first 200 bp of the transcriptional unit were considered; in the case of general and specific transcription factors as well as chromatin remodelers, reads were limited to the range -500 to +100 relative to the transcription start site (TSS).

### GO term and gene enrichment analysis

GO term and specific gene set enrichment analysis was performed using the GSEA tool (Mootha *et al*, 2003).

### Model for transcriptomic dynamics as a function of cell volume

We first defined four gene sets with high (*N*= 250), medium (*N*= 4500), low (*N*= 1000) and very low (*N*= 100) occupancy by RNA polymerase II as in experimental data shown in Fig 4D. Then, for each gene set (*i*), simple mass-action based equations were written in COPASI (Hoops *et al*, 2006):

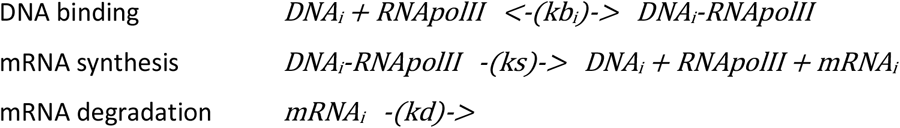

The range of nuclear volumes (2 to 40 fl, ca. 7% of cell volume) and the initial seed (1 μM) for fitting the RNA polymerase II concentration received realistic parameters corresponding to a wild-type haploid cell. Since synthesis (*ks*) and degradation (*kd*) constants did not have any effect on the relative change of mRNA levels with cell volume as predicted by the model, only the binding constant (*kb*) was set as variable for each gene set. The model was fitted to experimental data by the Levenberg–Marquardt method constraining binding constants from 10^7^ to 10^9^ M^−1^, which comprise the estimated binding constants of TFIIB (2×10^7^ M^−1^) (Lagrange *et al*, 1998) and TFIID (5×10^8^ M^−1^) (Hahn *et al*, 1989). Steady-state mRNA levels obtained from the four gene sets were first normalized to the total mRNA at every nuclear volume tested. Next, mRNA ratios were normalized to the average predicted from the corresponding gene set within the range of volumes analyzed. Since *ks* and *kd* are maintained constant, the predicted mRNA average was assumed linearly dependent on the binding constant (*kb*_*i*_) and gene number (*N*) for each gene set. Thus, conversion factors (*cf*_*i*_ = *kb*_*i*_ * *N*) for each bin were used with a general factor *gf* that makes *gf* * ∑ *cf*_*i*_ = 1 to normalize values as in the experimental datasets. Finally, to account for differences between experimental and predicted normalizing averages, variable factors

0.5 < *νf*_*i*_ < 2 were introduced for each gene set. Fits shown in Fig 4D were obtained with *vf*_*i*_ from 0.7 (very low occupancy) to 1.1 (high occupancy). Thus, the variable synthesis (*ks*) and degradation (*kd*) constants that likely affect the yeast transcriptome would account for deviations in normalizing averages smaller than 30%.

### Statistical analysis

Sample size is always indicated in the figure legend. Full datasets from single cells were plotted whenever possible but, when binned, bin size and statistics used are indicated in the figure legend. Pairwise comparisons were performed with a Mann-Whitney U test; and the resulting p-values are shown in the corresponding figure panels. The Spearman’s rank correlation test was used for non-parametric regression analysis.

## Data and code availability

RNA-seq fastq files are available as BioProject PRJNA879285, and the mathematical model from BioModels (ebi.ac.uk/biomodels) as MODEL2209080001.

## Acknowledgments

We thank E. Rebollo and M. Artés for technical assistance, D.F. Moreno for preliminary experiments, and B. Futcher for providing strains. We also thank C. Rose for editing the manuscript, and C. Gallego for helpful discussions. This work was funded by the Spanish Research Agency (PID2019-109638GB-I00) and the EU (FEDER). A.P.P. was supported by a FI fellowship (2019FI_B 00452) granted by AGAUR (Catalonia).

## Author contributions

P.J.V., A.P.P. and M.A. conceived and designed the study. P.J.V. and A.P.P. performed the experiments and analyzed the data. M.A. wrote and analyzed the mathematical model, and analyzed RNA-seq and ChIP-seq data. M.A. wrote the paper with input from P.J.V. and A.P.P.

## Dataset EV1

Transcriptomic summaries generated in this work.

---

**Figure EV1.**
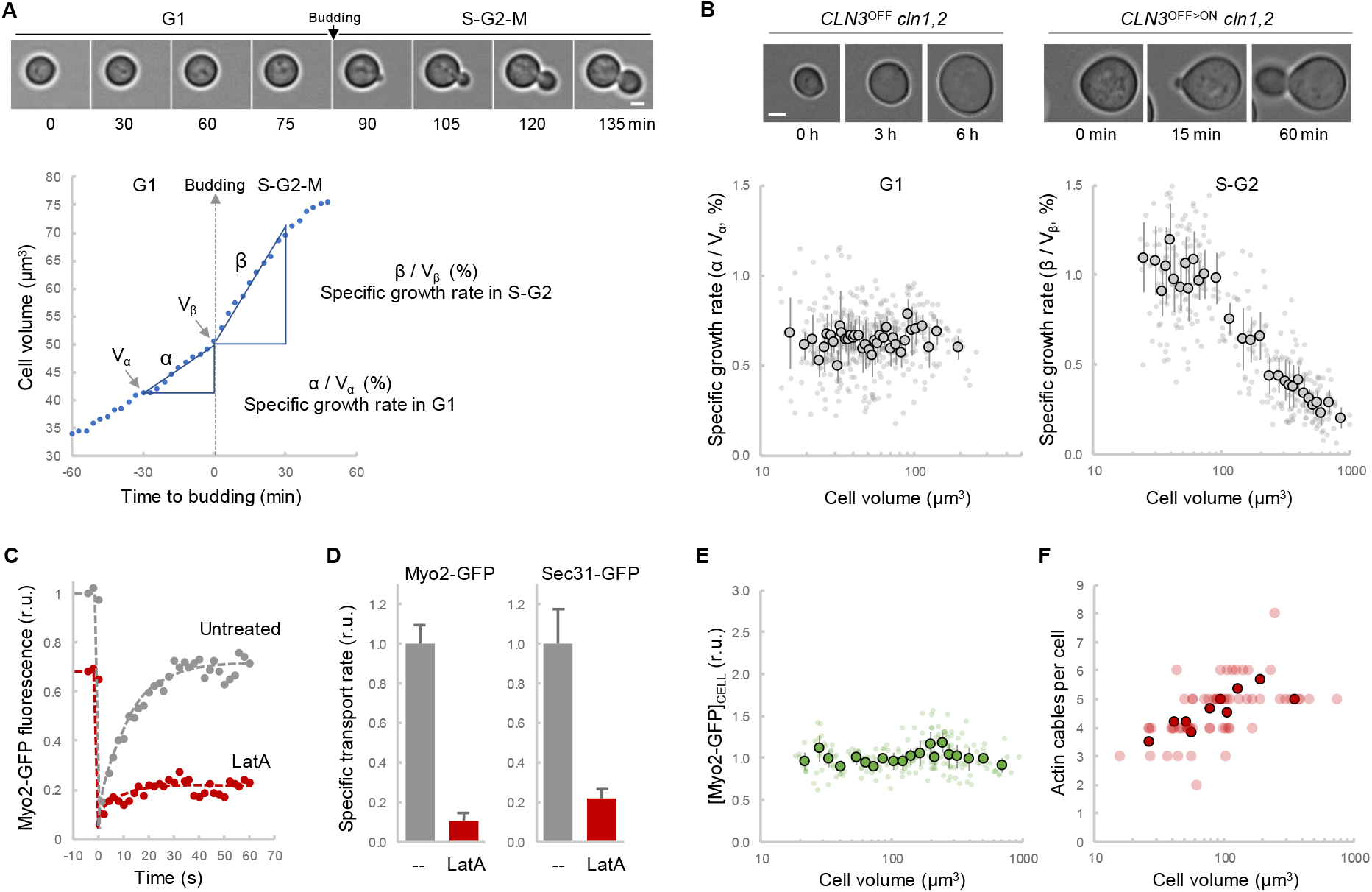
Growth potential in S-G2 phases is maximal around the critical size. A A representative yeast cell growing during the cell cycle. Scale bar, 2 µm. The plot shows cell volume as a function of time and the periods used to calculate the specific growth rates in G1 and S-G2. B Representative *GAL1p-CLN3 cln1,2* yeast cells arrested in G1 (*CLN3*^*OFF*^ *cln1,2*) in a glucose-based medium or entering the cell cycle after being transferred into a galactose-based medium to induce *CLN3* expression and trigger cell cycle entry (*CLN3*^*ON*^ *cln1,2*). Scale bar, 2 µm. Specific growth rates from G1 and S-G2 cells as a function of cell volume are plotted. Single-cell data (N > 300, small circles) are plotted. Mean values of binned data (bin size = 10, large circles) with confidence limits for the mean (α = 0.05) are also shown. C Representative Myo2-GFP kinetics during FRAP in cells treated or not with latrunculin A (LatA), an actin depolimerizing agent. D Specific transport rates of Myo2-GFP and Sec31-GFP to the bud in cells treated (N = 38) or not with latrunculin A (LatA, N = 24). Mean values per cell and confidence limits for the mean (α = 0.05) are plotted. E Myo2 concentration as a function of volume from cells as in Figs 1B and EV1B. Single-cell data (N = 200, small circles) are plotted. Mean values of binned data (bin size = 10, large circles) with confidence limits for the mean (α = 0.05) are also shown. F Number of actin cables as a function of cell volume from cells as in Figs 1B and EV1B expressing Abp140-GFP. Single-cell data (N = 61, small circles) are plotted. Mean values of binned data (bin size = 6, large circles) with confidence limits for the mean (α = 0.05) are also shown.

**Figure EV2.**
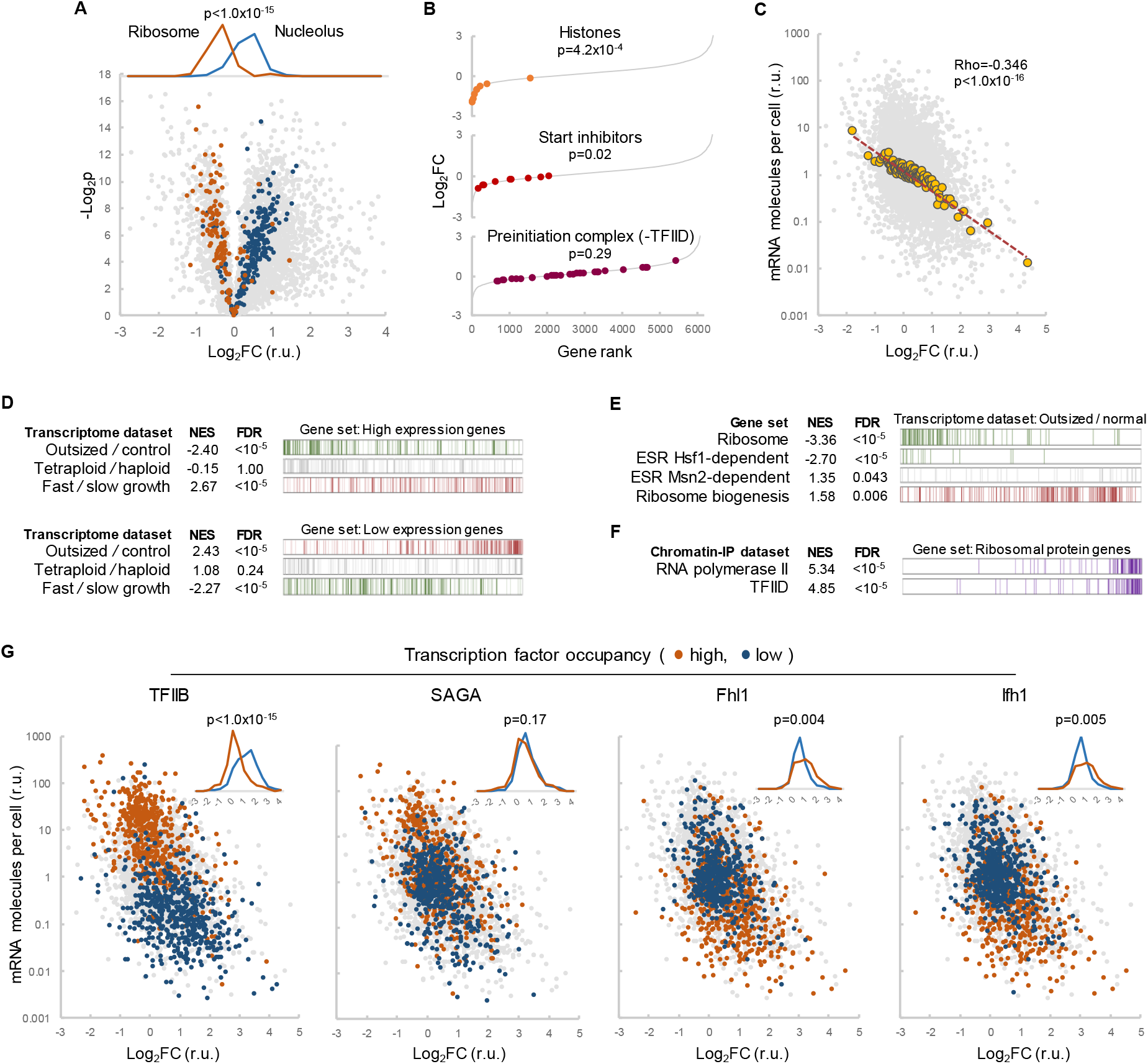
The transcriptome is extensively imbalanced in outsized cells. A Volcano plot of and expression in outsized *vs*. control cells highlighting genes coding for ribosomal (N = 137, orange) or nucleolar (N = 239, blue) proteins. Inset shows frequency distributions as a function of the expression fold change in outsized cells with the result of a Mann-Whitney U test. B Snake plots of gene expression in outsized *vs*. control cells highlighting genes coding for histones (N = 9), inhibitors of Start (N = 9) or PIC components excluding the TFIID complex (N = 30). The results of one-sample t tests are shown. C Plot of wild-type mRNA levels as a function of the expression fold change (Log_2_FC) in outsized cells. Single-gene (N = 6059, small dots) and binned (bin size = 50, large circles) are plotted with the corresponding regression line and the result of a Spearman’s rank correlation test. D-F Gene enrichment analysis of the indicated genesets and transcriptomic datasets. Barcode plots show the position of genes of the corresponding geneset within the indicated dataset ranked by the Log_2_FC. Normalized enrichment scores (NES) and FDR values are shown. G Plots of wild-type mRNA levels as a function of the expression fold change (Log_2_FC) in outsized cells highlighting genes with high (top 500 genes, orange) or low (bottom 500 genes, blue) occupancy of the indicated transcription factors in wild-type cells. Insets show the corresponding frequency distributions as a function of the expression fold change in outsized cells with the result of a Mann-Whitney U test.

**Figure EV3.**
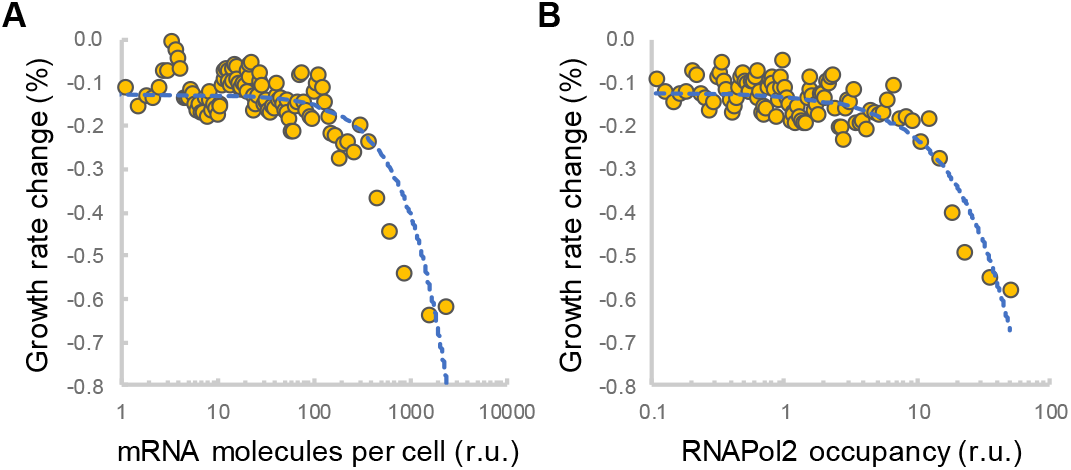
Cell size, growth competence and haploinsufficiency. A,B Growth rate change displayed by hemizygous strains of yeast genes (N = 5350) as a function of gene expression levels (A) or RNA polymerase II occupancy (B) in wild-type cells. The mean values of binned (bin size = 50) data and the corresponding regression lines are plotted.

**Figure EV4.**
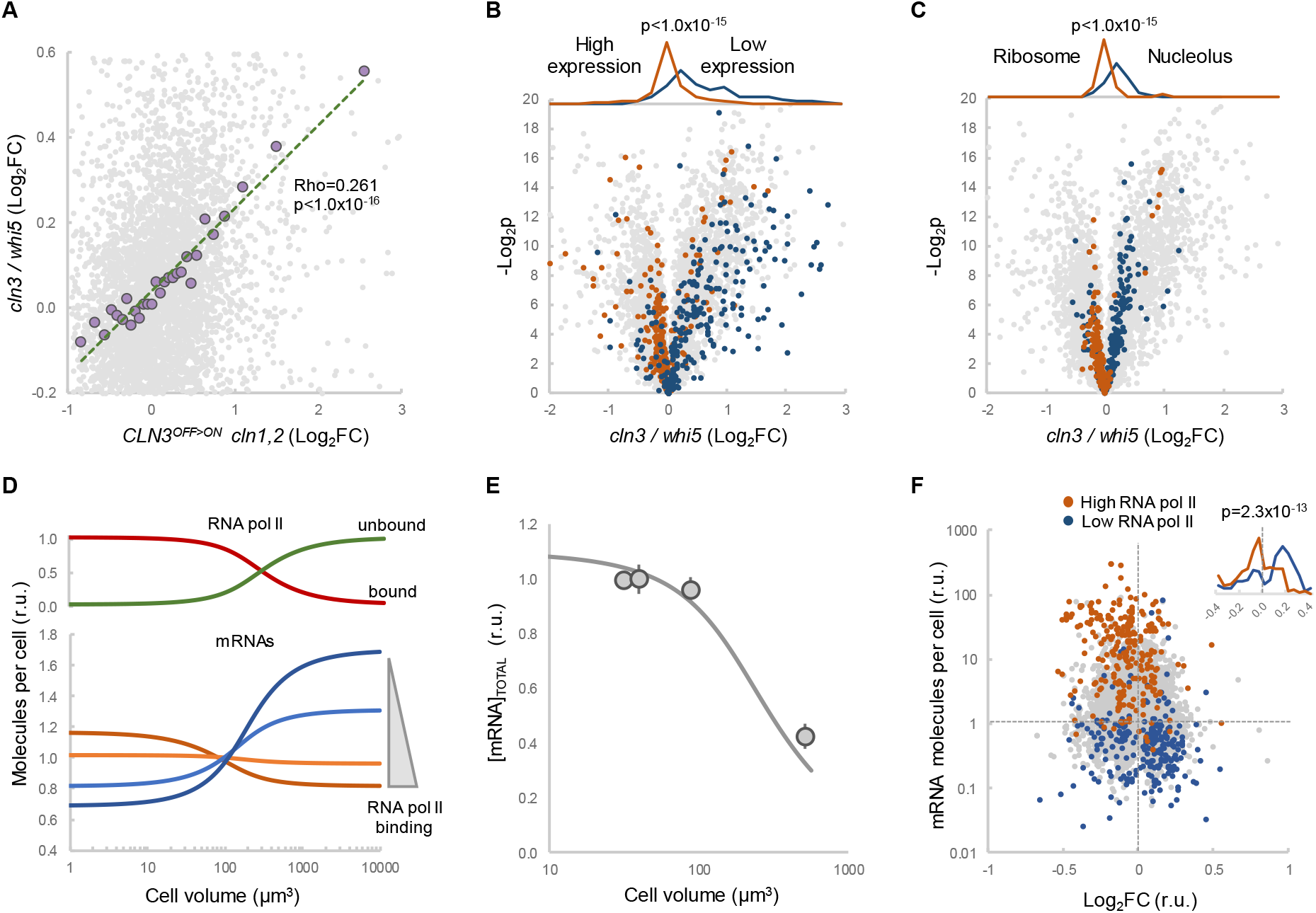
Inversion of the transcriptome by cell size and RNA polymerase II occupancy. A Fold change (Log_2_FC) values in large (*cln3*) *vs*. small (*whi5*) cells as a function of the fold change (Log_2_FC) in outsized (*CLN3*^OFF>ON^ *cln1,2*) *vs*. control cells. Single-gene (N = 5723, small dots) and binned (bin size = 200, large circles) data are plotted with the corresponding regression line and the result of a Spearman’s rank correlation test. B,C Volcano plots of differential expression in large (*cln3*) *vs*. small (*whi5*) cells highlighting genes with high (top 200 genes, orange) or low (bottom 200 genes, blue) expression levels in wild-type cells (B) or genes coding for ribosomal (N = 137, orange) or nucleolar (N = 239, blue) proteins (C). Insets show frequency distributions as a function of the expression fold change with the result of a Mann-Whitney U test. D,E Output of the model after being fitted to experimental datasets from yeast cells. Relative expression levels of genes with high (dark orange), medium (orange), low (blue) and very low (dark blue) affinity for RNA polymerase II (D, bottom), fraction of unbound (green) and DNA-bound (red) RNA polymerase II molecules (D, top), and predicted (E, thick line) and experimental (E, circles) total mRNA concentration as a function of cell volume. F Plot of wild-type mRNA levels as a function of the expression fold change (Log_2_FC) associated to HDAC inhibition highlighting genes with high (top 200 genes, orange) or low (bottom 200 genes, blue) RNA polymerase II occupancy in wild-type cells. Insets show the frequency distributions as a function of the expression fold change with the result of a Mann-Whitney U test.

**Table EV1.**
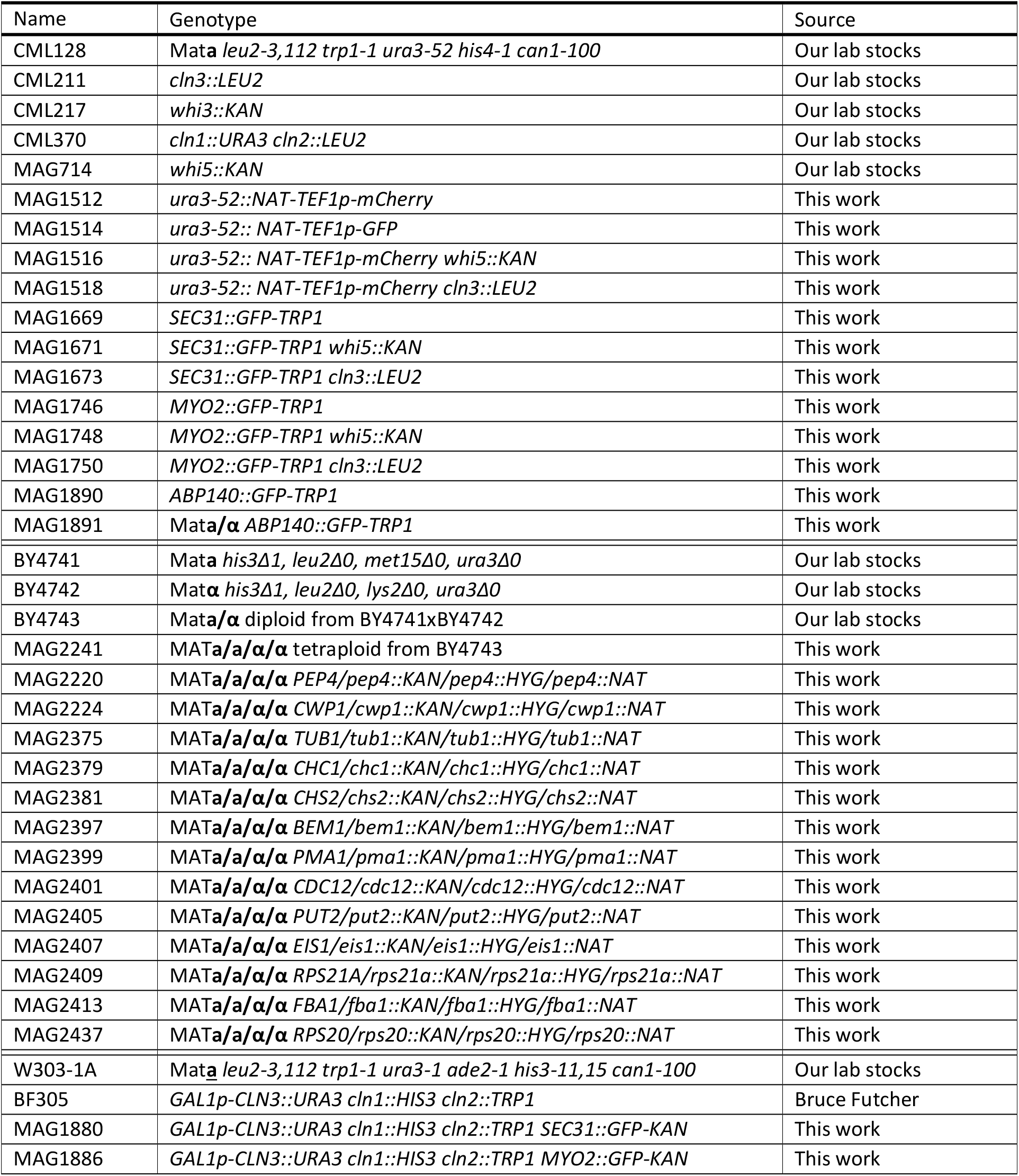
Yeast strains.

